# Analysis of Structural Variation Among Inbred Mouse Strains Identifies Genetic Factors for Autism-Related Traits

**DOI:** 10.1101/2021.02.18.431863

**Authors:** Ahmed Arslan, Zhuoqing Fang, Meiyue Wang, Zhuanfen Cheng, Boyoung Yoo, Gill Bejerano, Gary Peltz

## Abstract

The genomes of six inbred strains were analyzed using long read (LR) sequencing. The results revealed that structural variants (SV) were very abundant within the genome of inbred mouse strains (4.8 per gene), which indicates that they could impact genetic traits. Analysis of the relationship between SNP and SV alleles across 53 inbred strains indicated that we have a very limited ability to infer whether SV are present using short read sequence data, even when nearby SNP alleles are known. The benefit of having a more complete map of the pattern of genetic variation was demonstrated by identifying at least three genetic factors that could underlie the unique neuroanatomic and behavioral features of BTBR mice that resemble human Autism Spectrum Disorder (ASD). Similar to the genetic findings in human ASD cohorts, the identified BTBR-unique alleles are very rare, and they cause high impact changes in genes that play a role in neurodevelopment and brain function.

## Introduction

While commonly used next generation sequencing methods analyze ∼200-300 bp DNA segments (i.e., ‘*short read*’ (**SR**) sequencing) ^1^, recently developed ‘*long read*’ (**LR**) sequencing methods, which can analyze 20kb DNA segments ^2,3^, has enabled previously uncharacterized structural variants (**SV**) (i.e., genomic alterations >50 bp in size) to be evaluated. LR sequencing has been used to characterize genetic disease mechanisms that could not otherwise be analyzed ^1-3^; it has identified large genomic alterations in patients with Cardiac Myxomata (*PRKAR1A*) ^4^, Bardet–Biedl syndrome (*BBS9)* ^5^ and intellectual disability (*ARHGEF9*) ^6^. Mouse is the premier model organism for biomedical discovery, and many genetic factors affecting important biomedical traits have been identified from analysis of mouse genetic models ^7,8^. However, all murine genetic studies analyze SNP-based allelic variation, and the impact of SV has not previously been considered. As with human diseases, a more complete map of the pattern of genetic variation among inbred mouse strains that includes SVs could enable genetic discovery.

Therefore, LR sequencing was utilized to evaluate SVs in six selected inbred mouse strains, which included a strain (BTBR T+Itpr3tf/J, **BTBR**) that uniquely displays neuroanatomic abnormalities and behaviors characteristic of human Autism Spectral Disorder (**ASD**) ^9,10,11^. While a murine model cannot reproduce all aspects of a human neurobehavioral disease, the BTBR abnormalities reflect three important features of ASD: (i) neuroanatomic changes that include the complete absence of a corpus callosum (CC); (ii) a deficiency in engaging in social tasks; and (iii) abnormal repetitive behaviors ^11,12^. Despite the multiple studies that have been performed to date - which have employed epigenetic ^13^, genetic ^14^, transcriptomic ^15,16,17^ and proteomic ^15,18^ methodologies - the genetic basis for the BTBR neuroanatomic and behavioral abnormalities is not known. Analysis of the LR sequence for six strains and of SR sequence for 53 strains revealed that SVs are abundant in the genome of inbred mouse strains. Therefore, we investigated whether having a more complete map of the pattern of genetic variation in the BTBR genome, could identify genetic factors (SVs, Indels, SNPs) that have a high probability of contributing to the BTBR behavioral and neuroanatomic abnormalities.

## Results

### A genome-wide assessment of SV among six inbred mouse strains

LR genomic sequencing of six inbred mouse strains (BTBR, 129Sv1, C57BL/6, Balb/c, A/J, SJL) was performed, which had an average read length of 15.6 kb and >40x fold genome coverage (**Table S1**). The LR sequences were aligned to the reference C57BL/6 sequence; and SVs were identified that ranged in size from 50 bp to 10 kb. There were 48292, 48372, 41528, 41415, 5482, 45148 SVs identified within the 129Sv1, AJ, BALB, BTBR, C57BL/6, and SJL genomes, respectively (**Fig. 1A**). Since C57BL/6 is the reference sequence, only a very small number of SV were identified in its genome and relatively few passed subsequent quality control parameters, C57BL/6 SVs were not further analyzed here. For the five other strains, deletions and insertions were the most common type of SV, but duplications and inversions were also present. About 80% of the inversions (median 1551 bp) and 85% of the duplications (median 1695 bp) are > 500 bp in size; while 70% of the deletions (median size 209 bp) and 86% of the insertions (median size 156 bp) are < 500 bp; and 99% of the deletions and insertions are < 10 kb in size. Although duplications and inversions are much rarer than deletions, they are more common among the SV that are >10 kb in size. (**Figs. 1A-B**). Most (99%) SVs were within non-coding regions (intergenic, intronic, upstream, downstream, or regulatory), and were predicted to have a minor impact; but 628 SVs were predicted to have a major impact by causing stop- or start-codon loss, transcript ablation (most common) or amplification, or a frameshift (**Figs. 1C-D**). Since strain-specific SV alleles could affect the properties uniquely exhibited by an inbred strain, we identified 9032, 5648, 8537, 6018, and 3497 SVs that were uniquely present in the 129S1, AJ, SJL, BTBR, and BALB/c, genomes, respectively. Only 9.9% of the SVs are commonly shared by all 5 strains **(Fig. 1E)**. Since it has been suggested that repetitive elements, which include transposons, contribute to the generation of SV ^19^; it was of interest to find that 96% of deletions contained at least 1 repetitive element, and LINE1 and ERVK retrotransposon elements were among the most frequent type of repeat elements.

**Figure 1.**
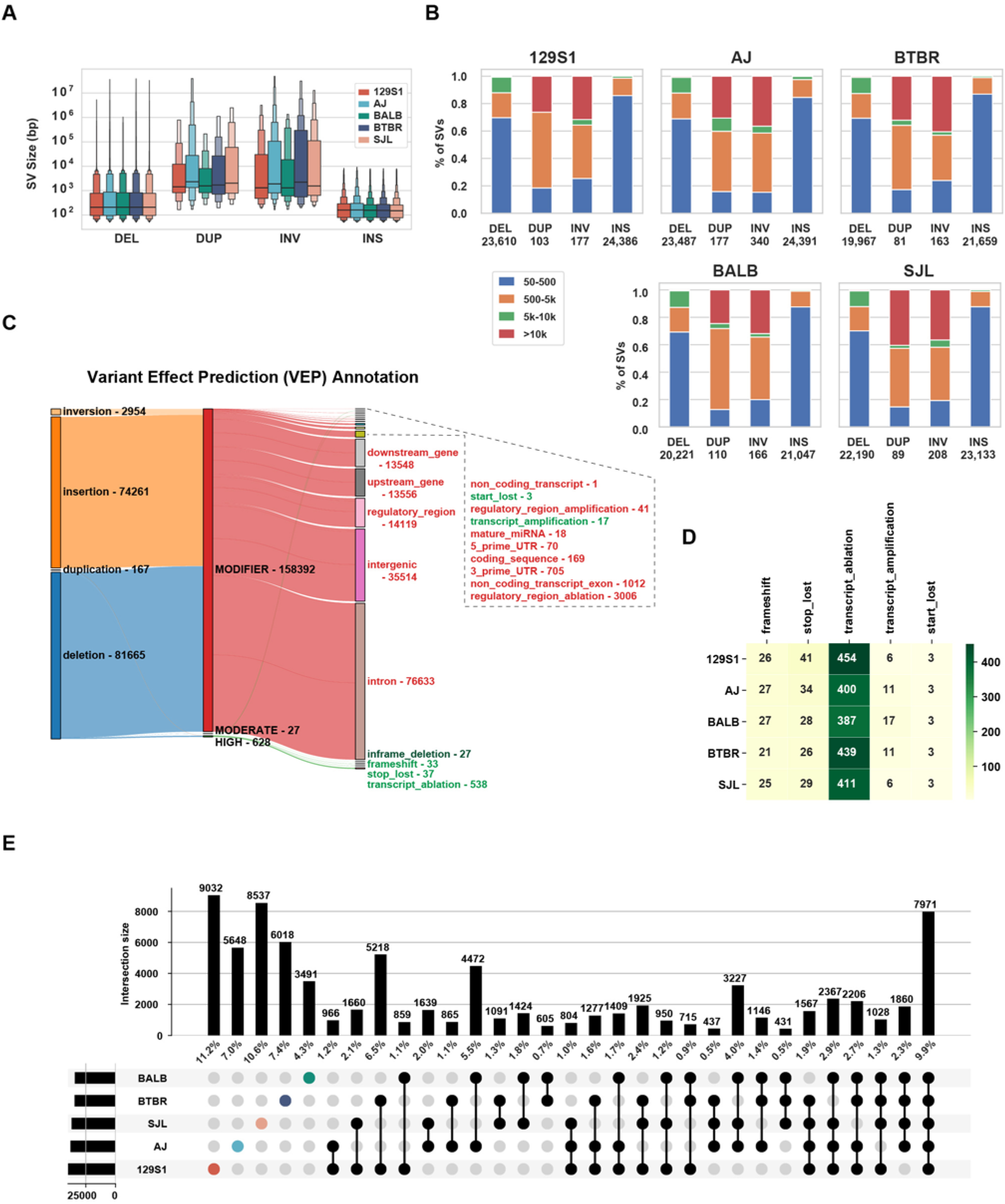
Characterization of SVs within the 129Sv1, A/J, SJL, Balb/c and BTBR genomes. **(A)** Letter-value boxplots ^64^ show the size distribution of the 4 different types of SVs present in the genomes of the 5 strains: DEL, deletions; DUP, duplications; INV, inversions; and INS, insertions. The wide box shows the 25-75% values, while each of the smaller boxes show 12.5% of each data set. (**B**) Each of the four types of SV are categorized according to their size for each strain, and the total number of each type of SV is shown at the bottom. **(C)** This Sankey diagram shows the predicted functional consequences for the four different types of SV, which are categorized by their estimated severity (MODIFIER, MODERATE, HIGH). Only 628 SV have a high functional impact (green), while the vast majority of SVs have a predicted minor impact. The number of SV with each type of functional annotation is indicated. (**D**). The number and type of the high impact SV present in each of the 5 strains are shown. (**E**) This UpSet plot shows the unique and shared SVs for each of the 5 strains. In the top graph, each vertical bar represents the number (and percentage) of SVs present in the strain(s) indicated in the intersection matrix below. In the intersection matrix, the total number of SVs in each of the 5 strains is indicated by the horizontal bar on the left; each colored dot indicates a single strain; and bars with 2 or more black dots indicate the number of shared SV among the strains indicated by the black dots.

### Comparing SVs identified by LR and SR sequence analysis

A previously described work flow ^20^ was used to analyze the available SR genomic sequence for 53 inbred mouse strains (average fold genome coverage 41x, range 19x to 168x) ^21^. This analysis identified 133,091 deletions, 11,162 duplications, and 1,608 inversions within their genomes (**Figs. 2A-B**). Several important observations emerged from analysis of these SVs (referred to as SR-SV). (i) Although the median size of the duplicated regions was 1162 bp, 32% of them were >10 kb; (ii) deletions are a major contributor to inter-strain differences in SV alleles, since they were 12-fold more abundant than duplications; and (iii) a large percentage of the deletions were strain-specific (**Fig. 2C**). To assess the quality of the SR-SV identified using the SR sequence workflow, we assessed their overlap with SV identified by LR sequence analysis for the 5 strains with available LR sequence. SR-SVs were only 25% of those identified by LR sequence analysis. Over 85% of SR-SVs overlapped with those identified by LR analysis and the vast majority (99%) of deletions were similarly classified by SR and LR analyses (**Fig. 3A**). However, there were significant differences in the results produced by the two analysis methods. Only 4.7 % of the duplications and 60% of the inversions identified by SR analysis were similarly classified by the LR analysis (**Fig. 3B**). Since the SR SV pipeline ^20^ has difficulty identifying insertions, and 32% of the SR duplications are > 10 kb in size (Fig. 3B), it is not surprising that many duplications (53-63%) identified by the SR sequence analysis were reclassified as insertions by the LR analysis (**Fig. S1**). These results indicate that LR sequencing enables many more SV to be detected, and that SV were more accurately classified using LR sequence.

**Figure 2.**
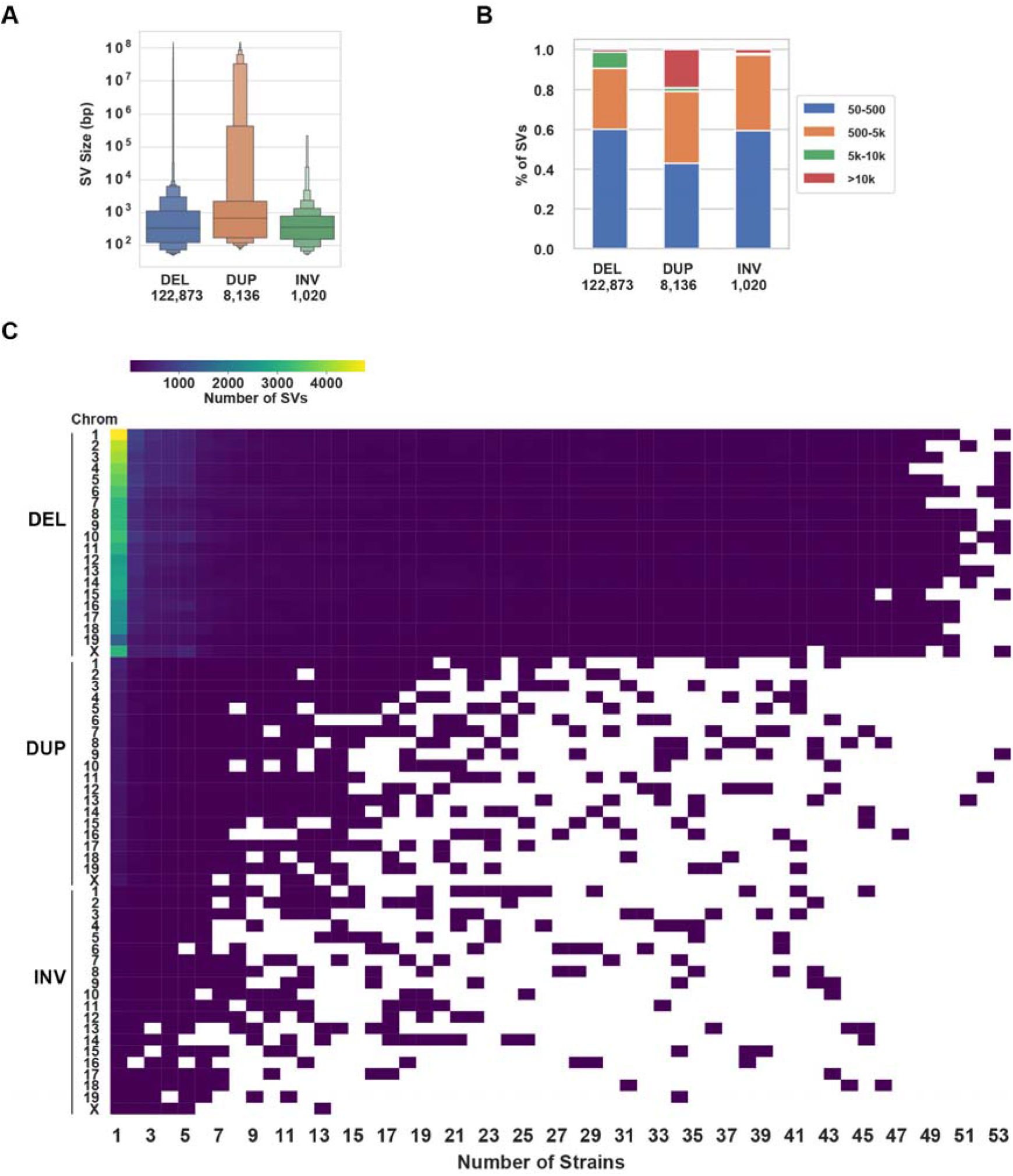
SV within the genome of 53 inbred mouse strains. (**A**) Letter-value boxplots show the size distribution of deletions, duplications and inversions, which have a median length of 337, 680, and 362 bp, respectively. The total number of each type of SV is shown at the bottom. (**B**) The SVs are categorized into 4 subgroups according to their size: 50-500 bp, 500 bp-5kb, 5-10kb, and >10kb. Over 90% of the deletions are <5 kb, 97% of the inversions are <5 kb, but 19% of the duplications are >10 kb. (**C**) The number of SVs are categorized according to their type and chromosomal location, and by the number of inbred strains with a strain-shared SV. Each box color indicates the number of each type of SV according to the scale shown at the top. A white area indicates that shared SVs were not found for that number of strains. Deletions are the most common type of SV, and the majority are uniquely present in one strain.

**Figure 3.**
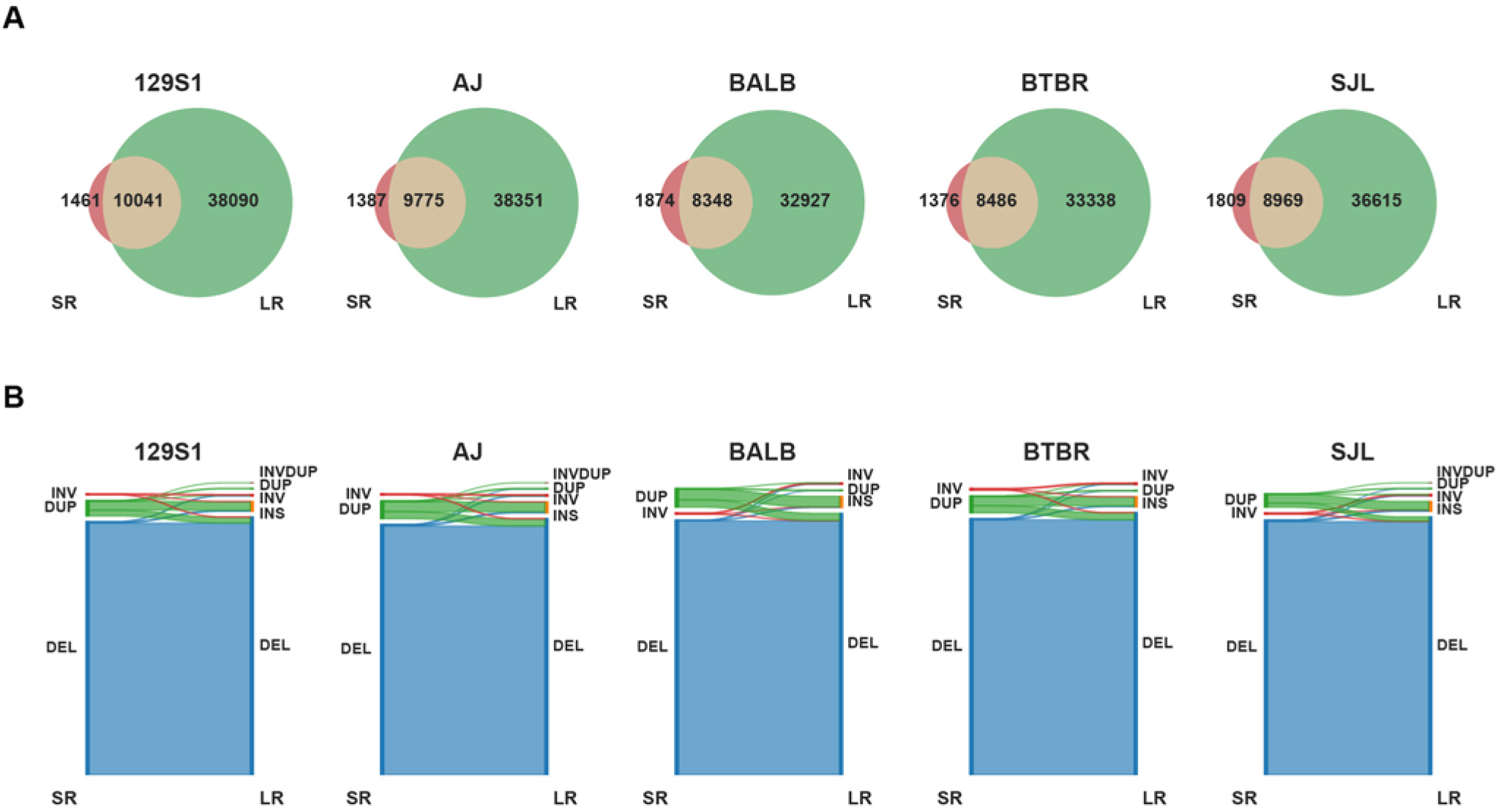
SR and LR SV comparison. (**A**) Venn diagrams show the overlap of SV identified by LR or SR sequence analysis for 5 strains. (**B**) Sankey diagrams indicate the number and type of SR-SVs that were confirmed after analysis of the LR sequence for each strain. Overall, the percentage of SR-SVs that were confirmed by the LR analysis are: 99.4% for DEL, 5% for DUP, and 61.3% for INV. Duplications >10 kB was the source of the largest discordance between the SR and LR results.

### The relationship between SV and SNP allele

We examined the relationship between the 146K SR-SV and the previously identified 22M SNP alleles in the genomes of 53 inbred strains ^21^. Just as for a SNP allele, the presence or absence of a SV was treated as an individual allelic variant, even though it impacts >50 bp. The average distance for a 50% decay in the mean linkage disequilibrium (LD) (*r*^2^) calculated using both SNP and SV alleles (30-38 kb) was similar to that when only SNP alleles were used (31-40 kb) (**Fig. S2**). We then examined the relationship between SV and SNP alleles by characterizing LD within localized (±50 kb) genomic regions; and the strain-unique and shared (present in ≥2 strains) SR-SVs were separately analyzed. Across all 53 inbred strains, only 3% of strain-unique SVs (1956) are in complete LD with nearby SNP alleles (**Tables, S2, S3A**), while 41% (32748) shared SVs are in complete LD with nearby SNPs. We also analyzed the subset of SVs located within the 21,832 protein-coding genes. Across the 53 inbred strains, a protein coding gene had an average of 4.8 SVs, and a SV is in complete LD (*r*^2^ = 1) with an average of 4.5 nearby SNPs. Thus, irrespective of whether whole genome or intergenic regions were analyzed, the SV-SNP allelic relationships are quite similar (**Table S3B**).

Our prior analysis of genome-wide SNP allele relationships separated the 53 inbred strains into four sub-groups ^22^. The six sub-group 1 strains are derived from a C57BL ancestor; sub-groups 2 (17 strains) and 3 (25 strains) contain most of the classical inbred strains; and the five sub-group 4 strains are wild-derived (**Table S2**). The classical inbred strains in sub-groups 1-3 have 2.1-fold more shared SVs than strain-unique SVs, which is probably due to their having shared genomic segments derived from ∼four ancestral founders ^23,24^ (**Table S2**); while the wild-derived strains have 2.3-fold more strain-unique than shared SVs, which is consistent with their increased genetic divergence. The number of SVs uniquely present in the genomes of the five wild-derived strains is 59% of the total number of SVs identified in all 53 strains. As discussed in supplemental note 1, inclusion of the wild-derived (group 4) strains dramatically reduced the LD relationships between SV and SNP alleles among the group 1-3 strains (**Fig. S3)**.

### Identification of three candidate genetic factors for the ASD-like properties of BTBR mice

Genetic studies have identified multiple BTBR genomic regions that make distinct contributions to their ASD-like abnormalities ^14^. Since we now have BTBR LR genomic sequence, along with SR genomic sequence for the 52 other inbred strains ^21^, we investigated whether BTBR-specific genetic factors that contribute to these abnormalities could be identified. Since ASD patients have a much higher frequency of highly disruptive mutations (copy number variation ^25^, premature termination codons (PTC), frameshift) within neuro-developmentally important genes that are very rare in the general population ^26,27^; we sequentially identified BTBR-specific SV and SNP alleles that have a high impact on genes expressed in brain. This analysis of SV also included high impact indels (i.e. SV ≤ 50 bp in size). We analyzed the LR BTBR sequence and identified 8 BTBR-unique SV that could significantly alter transcripts (**Table S4**), but none had a previously reported connection with ASD. We then identified 6 BTBR-unique high impact indels (**Table 1**). An 8-bp frameshift deletion at the end of exon 2 of the *dorsal repulsive axon guidance protein* (*Draxin)* was of interest. *Draxin* is located within an interval that contributes to multiple BTBR commissural abnormalities ^14^. The truncated BTBR Draxin protein is missing key binding domains that are essential for its function in regulating neurite outgrowth (**Fig. 4)**. This makes BTBR mice the equivalent of *Draxin* knockout (KO) mice, which have abnormal development of the CC and forebrain commissures ^28^. Developing CC axons are guided to their final destination by midline glial structures ^29^ that are absent in *Draxin* KO mice ^28^. Human GWAS have identified an association between *DRAXIN* alleles and susceptibility to schizophrenia (rs4846033, p-value 4×10^−6^, intergenic variant) ^30^ and ASD (rs12045323, 7×10^−6^, intronic variant) ^31^. However, the CC defects in *Draxin* KO mice were quite variable ^28,32^, while the CC is completely absent in BTBR mice, which indicates that other BTBR genetic factors contribute to this defect. To identify them, we analyzed 29 SNPs that altered the sequence of the encoded protein (**cSNPs**) with BTBR-specific alleles in the 28 genes expressed in brain (see supplemental note 2, **Table 2**). Only one introduced a PTC in C1q tumor necrosis factor related protein 5 (C1QTNF5, which is also referred to as *Ctrp5*), which truncated the Ctrp5 protein at position 173 (*Q173X*) within its C1q domain (**Fig. 5A**). *Ctrp5* belongs to a superfamily of proteins that play diverse roles in mammalian physiology ^33^, and it is expressed in astrocytes ^34^. Its C1q domain is essential for multimerization ^35^ and for its binding to a serine protease (High temperature requirement A1, **HTRA1**) ^36,37^ that degrades extracellular matrix (ECM) proteins ^38,39^. HTRA1 protease activity is stimulated by interaction with the CTRP5 COOH-terminal region (**Fig. 5B**) ^40^. HTRA1 degrades amyloid precursor protein **(APP)** ^38^, other Alzheimer’s Disease-related proteins (Apo4, tau), and proteins that affect the ECM (ADAM9, Clusterin, C3, vitronectin) ^39^. Since BTBR Ctrp5 has a reduced ability to stimulate HTRA1 proteolytic activity, this could alter ECM processing and astrocyte development. Several human *CTRP5* mutations ^41^ are associated with late-onset retinal degeneration (**L-ORD**) ^40^. A C57BL/6 mouse with a knockin (**KI**) of a L-ORD causing human mutation (*S163R*) ^42^ developed retinal pathology ^40,43^, while the 129Sv *S163R* KI mouse did not ^42^. Hence, another genetic factor is required for the appearance of *Ctrp5* mutation-induced retinal defects. By this mechanism, the *Draxin* and *Ctrp5* mutations could jointly contribute to the complete agenesis of the BTBR CC. Another BTBR-specific 26 bp frameshift deletion within *Parp10* was also identified (Supplemental note 3, **Fig. S4**), but the role of Parp10 in ASD or CC formation has not yet been fully established.

**Table 1.**
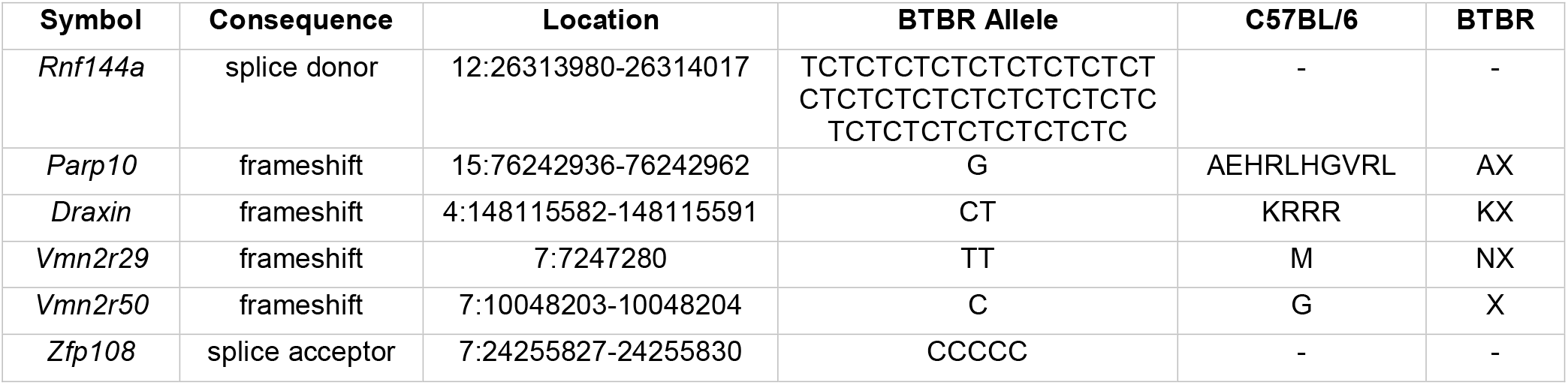
BTBR-unique high impact indels. The gene name, associated transcript; functional consequence, location, BTBR unique allele, and amino acid change(s) (C57BL/6 vs BTBR) for 6 high impact BTBR-unique indels are shown.

**Table 2.** Genes with BTBR-unique cSNPs that are expressed in brain. The gene name, the amino acid position (POS), and the amino acid in the reference (C57BL/6, REF) and BTBR-unique alleles are shown for the 29 cSNPs within the 28 genes that are expressed in brain.

|  | **Gene** | **Pos** | **Ref** | **BTBR** |
| --- | --- | --- | --- | --- |
| 1 | *Ctrp5* | 173 | Q | X |
| 2 | *Washc5* | 176 | R | W |
| 3 | *Tnrc18* | 327 | R | L |
| 4 | *Pkp4* | 335 | S | L |
| 5 | *Psd3* | 302 | V | E |
| 6 | *Gtf3c1* | 1708 | A | V |
| 7 | *Scn1a* | 66 | G | R |
| 8 | *Sorbs3* | 431 | A | T |
| 9 | *Akap6* | 2253 | T | M |
| 10 | *Nuak2* | 327 | P | S |
| 11 | *Zfp622* | 177 | P | S |
| 12 | *Asphd1* | 34 | A | T |
| 13 | *Ankrd17* | 962 | T | M |
| 14 | *Macf1* | 2335 | T | K |
| 15 | *Ptprc* | 41 | H | Q |
| 16 | *Tnc* | 1552 | T | S |
| 17 | *Tnc* | 1559 | V | I |
| 18 | *Ehbp1* | 855 | R | Q |
| 19 | *Tmem43* | 240 | R | Q |
| 20 | *Ppp1r16b* | 223 | D | N |
| 21 | *Ctbp2* | 400 | A | S |
| 22 | *Polrmt* | 1003 | S | N |
| 23 | *Sacs* | 1029 | N | D |
| 24 | *Bin3* | 116 | S | T |
| 25 | *Mical3* | 1165 | V | M |
| 26 | *Myo9a* | 810 | R | K |
| 27 | *Tm2d2* | 211 | T | P |
| 28 | *Oxct1* | 50 | V | I |
| 29 | *Col27a1* | 149 | V | I |

**Figure 4.**
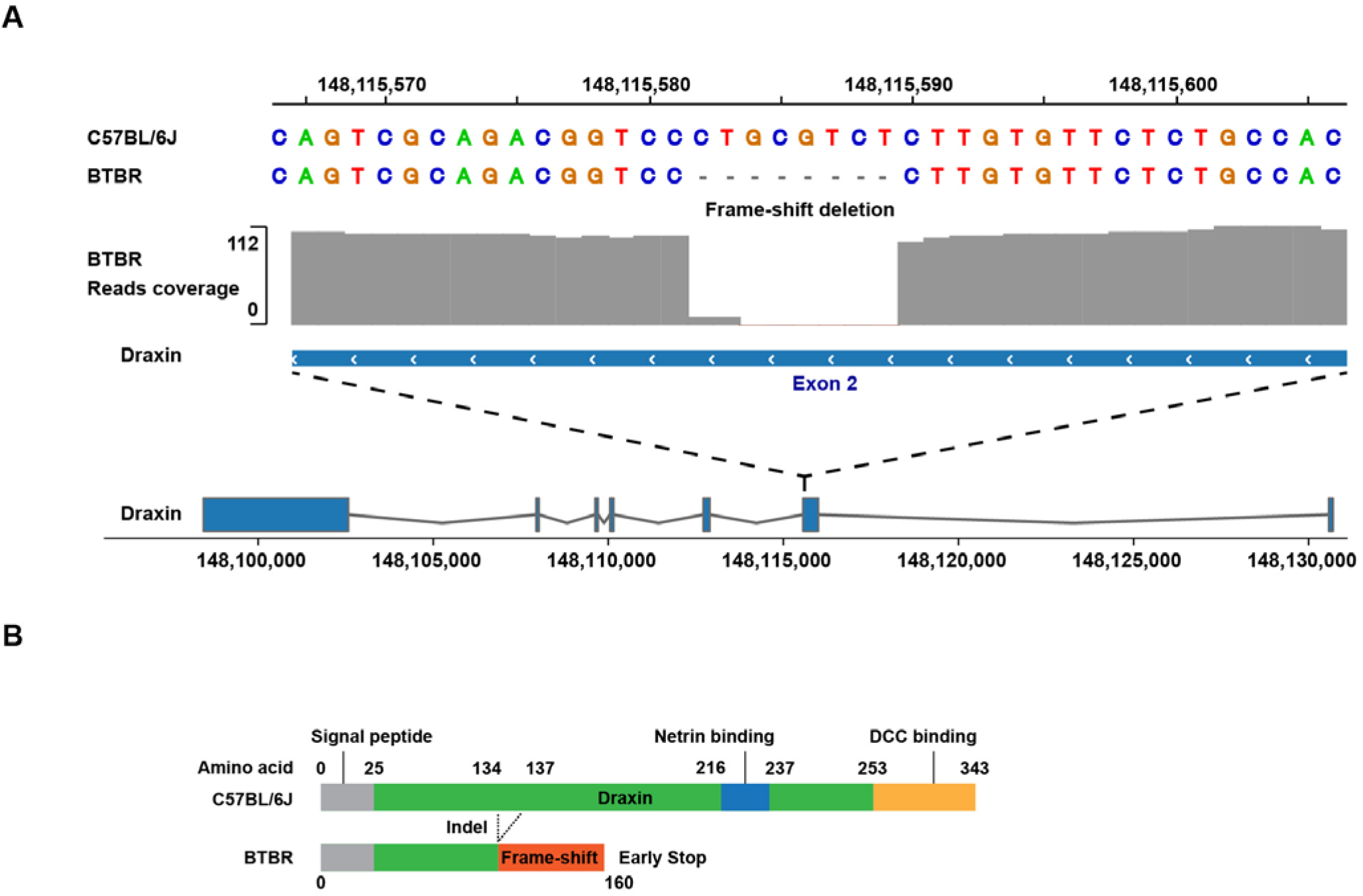
BTBR mice produce a non-functional Draxin protein. **A**) BTBR LR genomic sequence analysis identified an 8 bp deletion at the 3’ end of exon 2 of *Draxin*, which is not present in 52 other strains with available genomic sequence. **B**) While the full length draxin protein has 343 amino acids; the 8 bp frameshift deletion (that occurs between amino acids 134-137) generates a termination codon at amino acid 160 of the BTBR Draxin protein, which lacks the Netrin and DCC binding domains that are essential for its function in neurodevelopment.

**Figure 5.**
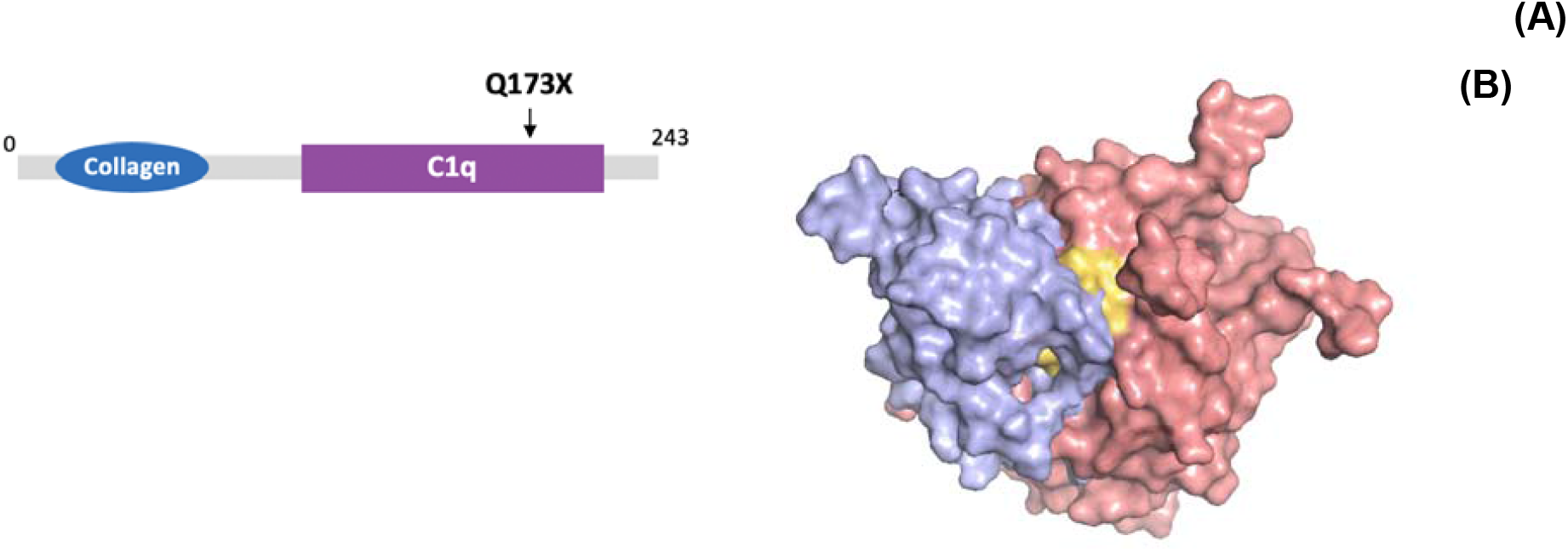
BTBR mice produce a truncated Ctrp5 protein. (***A***) A BTBR-unique SNP allele (*Q173X*) introduces a premature termination codon that truncates the 243 amino acid Ctrp5 protein at position 173 within its C1q domain. (**B**) A 3D reconstruction from ^40^ of the interaction of a trimeric complex of CTRP5 (red) with the PDZ domain of HTRA1 (blue). The COOH-terminal peptide sequence in CTRP5 (yellow), which is eliminated by the BTBR 173X PTC, binds to the PDZ domain of HTRA1.

To explain the behavioral phenotype, we searched PubMed papers to examine the relationship between the 28 genes in Table 2 and ASD; *Scn1a* had the strongest relationship with autism (**Fig. 6A**). *Scn1a* encodes the pore-forming α-subunit of the voltage-gated sodium channel type-1 (Nav1.1) ^44,45,46^. A BTBR-specific SNP allele (*Gly66Arg*) within its cytoplasmic domain disrupted a highly conserved tyrosine-based internalization and sorting signal (65-Y**G**DI-68) ^47^ that is present in many membrane proteins that traffic to the cell membrane ^47,48^, which mediates the interaction between Scn1a and the μ subunit of the AP-2 complex (Ap2m1) (**Fig. 6B**). The charged and bulkier BTBR *Arg66* allele could reduce this interaction, which is facilitated by the absence of a side chain in the *Gly66* residue (Fig. 6B). The amino acid sequences of murine Scn1a and human SCN1A are 98% identical; and both contain the YGDI sequence. Since there are no human cSNPs at the *Gly66* site in the human population ^49^, we examined the overall tolerance of human *SCN1A* to allele-induced loss of function (**LoF**) using a program (LoFtool) that was optimized for assessing gene tolerance to functional variation ^50^. The results (1.38 x 10^−04^) indicate that *SCN1A* is highly intolerant of allelic variation. It is also possible to speculate that the recognition motif in SCN1A could have a role in cognitive function and mood; multiple antidepressants with chemically distinct structures decreased the YXXØ-AP2M1 interaction ^51^. Also, since mutations in *SCN1A* ^46^ or *AP2M1* ^52^ are independently associated with ASD susceptibility, a BTBR-unique allele that alters their interaction is highly likely to contribute to its ASD-like behaviors.

**Figure 6.**
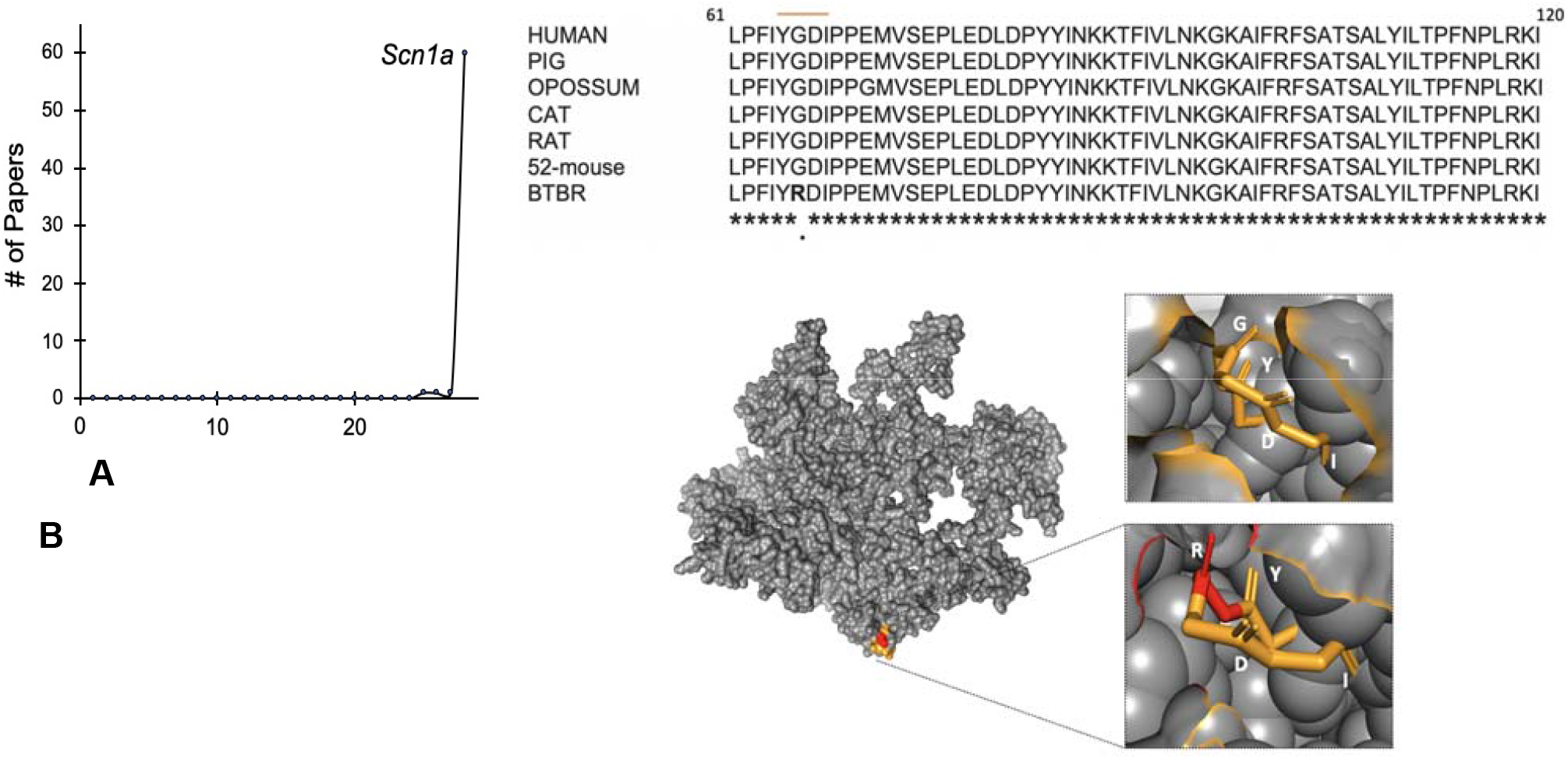
The effect of a BTBR-unique *Scn1a ARG66* allele. (**A**) Graphical representation of the number of PubMed papers that co-mention ASD and each of the 28 genes with a BTBR-unique cSNP. From this analysis, *Snca1* was most strongly associated with ASD (n=60 papers); while only 3 genes (*Tnc, Pkp4, Akap6*) had a single ASD-associated paper; and the 24 other genes did not have any ASD-associated papers. (**B**) *Top*: An alignment of the Scn1a cytoplasmic domain region containing the YXXØ motif (positions 65-68) for six mammalian species is shown. The nonpolar Gly66 within the YGDI motif is conserved in all species and in 52 inbred mouse strains; while the *Arg66* allele, which produces a major amino-acid change with a Blossom-62 score of -2, is only present in BTBR mice. *Bottom*: A murine Scn1a (ID: A2APX8) homology model showing the location of the surface exposed YXXØ motif in the cytoplasmic domain (shown in color), and enlarged images showing the conserved Y**G**DI motif (upper panel, gold) and the YRDI motif is shown in BTBR mice (lower panel; red for Arg66 and gold for the unchanged amino acids).

## Discussion

Several important features about the impact of SVs on the pattern of genetic variation in the genome of inbred strains are revealed by our analysis. (i) SV are quite abundant (average 4.8 per gene), which indicates that they are highly likely to impact genetic traits. (ii) SV and SNP alleles are more concordant among the classical inbred strains than among wild-derived strains. (iii) LR sequencing of additional strains is needed to produce a more complete map of the pattern of genetic variation among inbred mouse strains. As discussed in supplemental note 4, our ability to infer whether a known SV is present from analysis of nearby SNP alleles is quite limited, and it is very difficult to detect or characterize strain-specific SV using SR sequence.

Previous analyses of BTBR mice have identified many genes and potential pathways that could contribute its ASD-like properties ^13,14,15,16,18,53,54^. However, because all of the prior studies did not have access to the complete BTBR genomic sequence, they could not generate specific hypotheses about the genetic basis for its ASD-like features. By analyzing SNPs, indels and SV in the BTBR genome, we identified at least three BTBR-unique alleles that are likely to contribute to its ASD-like features because they cause major changes in genes with a role in neurodevelopment. Two cause protein truncating mutations; and one (*Scn1a Gly66ARG*) occurs within a highly conserved region in a gene that is intolerant of allele-induced LoF. Also, human *SCN1A* mutations cause Dravet’s syndrome (DS), which is a developmental epilepsy syndrome that is accompanied by ASD-related neuropsychiatric comorbidities ^55,56^. *Scn1a* haploinsufficient mice (*Scn1a*^+/-^) exhibited hyperactivity and some features of ASD, which include stereotyped behaviors and social interaction deficits ^57^. Like the retinal pathology in L-ORD KI mice, the *Scn1a*^+/-^ effects are strain dependent: seizures only occurred in *Scn1a*^+/-^ mice with a C57BL/6 (but not a 129) background ^58-61^ and were also dependent on the type of interneuron with the conditional KO ^62^.

The identified BTBR candidate factors are consistent with the results obtained from several large human cohorts; ASD patients have a much higher frequency of disruptive mutations in neurodevelopmentally important genes, which are very rare in the general population ^26,27^. Our results are also consistent with those obtained from analysis of BTBR intercross progeny. We identified BTBR-specific alleles that could separately impact their neuroanatomic and behavioral abnormalities, including one (*Draxin*) within a chromosomal region contributing to its commissural abnormalities ^14^. While their impact must be experimentally confirmed, these factors provide a starting point for understanding how a set of genetic changes, each of which is very rare in the mouse population, jointly produce an ASD-like phenotype. The fact that a human ASD-related phenotype in mice has an oligogenic basis is not unexpected. For example, while trying to produce a murine model of a human congenital cardiomyopathy, cardiac abnormalities did not appear until three different causative alleles – each homologous to a human disease-causing allele - were CRISPR engineered into three different murine genes ^63^. The cardiac abnormalities were not observed in mice with engineered changes in one or even two genes. Of importance, CRISPR methodology enables us to genetically engineer changes into the genome of any inbred strain, and tissue can be readily obtained for analysis at various developmental stages. Therefore, subsequent analyses of the effect that these BTBR-unique genetic factors have on neurodevelopment and behavior in mice should provide new insight into the pathogenesis of ASD. More broadly, these results demonstrate how obtaining a more complete picture of the pattern of genetic variation among inbred mouse strains can facilitate genetic discovery.

## Competing interests

The authors declare that they have no competing interests.

## Data availability

Upon acceptance of this paper, the sequencing data will be made available in the Sequence Read Archive (SRA).

## Funding

This work was supported by a NIH/NIDA award (5U01DA04439902) to GP.

## Author contributions

G.P. and A.A. formulated the project with input from all authors. Z.C. and A.A. generated experimental data. A.A, M.W. B.Y. and Z.F. analyzed the data; G.B. helped with the analysis. B.Y. A.A. and Z.F. contributed code. G.P., Z.F. and A.A. wrote the paper with input from all authors.

## Abbreviations

ASD: Autism Spectral Disorder
CC: corpus collosum
DS: Dravet’s syndrome
ECM: extracellular matrix
KI: knockin mouse
KO: knockout mouse
L-ORD: late-onset retinal degeneration
LOF: loss of function
LR: long read
PTC: premature termination codons
SR: short read
SNP: single nucleotide polymorphism
SR-SV: structural variants identified using SR sequence
SV: structural variant.

## Notes

### Competing Interest Statement

The authors have declared no competing interest.

