## supplemental_info for "Analysis of Structural Variation Among Inbred Mouse Strains Identifies Genetic Factors for Autism-Related Traits"

*Supplemental note 1: The effect of strain subgroups.* Analysis of genome-wide SNP allele relationships separated the 53 inbred strains into four sub-groups ^1^. The six sub-group 1 strains are derived from a C57BL ancestor; sub-groups 2 (17 strains) and 3 (25 strains) contain most of the classical inbred strains; and the five sub-group 4 strains are wild-derived (**Table S2**). We identified 11443, 46323, 40244 and 114086 SR-SVs within the genomes of the 53 inbred strains in sub-groups 1-4, respectively (**Table S3A**). We examined the LD between the SR-SV and SNPs within strain sub-groups, and the LD relationships were quite dependent upon whether all 53 strains or strain subgroups are analyzed. Over 75% or 84% of the strain-unique SVs are in perfect LD with ≥1 nearby SNP when sub-group 1-3 stains and subgroup 4 strains are separately evaluated, respectively. Inclusion of the group 4 strain-unique SVs dramatically reduced the LD relationships among the group 1-3 strains. Within each of 4 sub-groups, >71% of shared SVs are in complete LD with nearby SNPs; while across all 53 inbred strains, about 41% (32748) are in complete LD with nearby SNPs. Among the subset of SVs within the 21,832 protein-coding genes, sub-group 1 strains have the lowest number (mean 2.1) of SVs per gene; sub-group 2 and 3 strains have 2.5 SVs per gene on average, and 2 of them are completely linked ($r^{2}=1$) with adjacent SNPs; and sub-group 4 strains have more SVs per gene (mean 4.1). **Fig. S3** illustrates how the relationship between SV and SNP alleles within a gene is affected by whether all strains or strain subgroups are analyzed.

*Supplemental note 2: BTBR-specific SNP alleles*. The genomic sequences of BTBR and 52 other inbred strains were analyzed to generate a 22M SNP database with alleles that cover 53 inbred strains ^2^. Since a BTBR-unique allele could contribute to its unique behavioral or neuroanatomic phenotypes, this SNP database was computationally analyzed to identify SNP-alleles that were only present within the BTBR genome. Of the 35,000 SNPs with BTBR-specific alleles, 138 SNP alleles altered the amino acid sequence of 90 different genes (**cSNPs**). The low number of cSNPs is consistent with our finding that cSNPs account for only 0.2 - 1.0% of all SNPs in the database. We then used the information in the Allen Institute Expression Atlas database ^3^ and found that 28 of these genes were expressed in brain. Thus, only 29 cSNPs (in 28 genes) were expressed in brain; and only one uniquely introduced a PTC into the BTBR genome (**Table 2**).

*Supplemental note 3: A Parp10 frameshift deletion*. We also identified BTBR-unique 26 bp frameshift deletion in exon 2 of *Parp10 (ARTD10),* which generates a termination codon after amino acid 66. The truncated BTBR protein lacks the catalytic and other domains of Parp10 that are essential for its function (**Fig. S4).** *Parp10,* which is located within an interval on chromosome 15 that contributes to BTBR commissural abnormalities ^4^, is a mono-ADP ribosylation (marylation) enzyme. Because of its many substrates, it is involved in transcriptional regulation, DNA damage repair, and other cellular processes ^5,6^. While a functional role for Parp10 in ASD has not been directly demonstrated, some evidence suggests that it could play a role. Through marylation of protein kinase C (PKC), Parp10 alters the activity of a voltage-gated potassium channel (Kcna1, Kv1.1), which regulates hippocampal neuron excitability ^7^; and Kv1.1 has been shown to modulate ASD-related behaviors in mice with another ion channel mutation ^8^. Also, a patient with a homozygous *PARP10* deficiency had neuro-developmental delay and defective DNA repair ^9^. PARP10 has been shown to regulate NF-κB ^10^ and Wnt signaling ^11^, which are pathways that are known to affect neurodevelopment.

*Supplemental note 4: Can SV be imputed from analysis of SR sequence?* Given that we now have LR sequence data available for only 5 strains, a major question is whether SVs can be characterized for the other inbred strains using SR sequence data? Our results indicate that it is possible to predict whether a known SV (i.e., one identified from analysis of LR sequence data) is present in other strains based upon analysis of nearby SNP alleles. However, since only 24% of SVs are in complete LD with nearby SNPs across the 53-strain panel, the level of certainty for SNP allele-based SV prediction is quite limited. Even when the analysis is confined to SV predictions within a strain subgroup, only 41% of shared SVs are in complete LD with nearby SNPs. Thus, the ability to impute whether SVs are present based upon nearby SNP alleles is limited, and this holds true even when the analysis is confined to the classical inbred strains. Our results also indicate that it can be difficult to identify sites with strain-unique SVs using only SR sequence data. Although >75% of the strain-unique SVs are linked with ≥ 1 adjacent SNP within each of the 4 strain subgroups, only 3% of strain-unique SVs have perfect LD with nearby SNP alleles in the 53-strain panel. The improvement of within group predictions results from the fact that each subgroup contains either closely related strains (subgroups 2 and 3) or has limited variation due to it having a small number of strains (subgroups 1 and 4). Nevertheless, the significant uncertainty associated with imputed SV limits their utility.

**Material and methods**

*LR DNA sequencing.* DNA was extracted from tails obtained from male 129Sv1, BTBR, SJL, A/J, Balb/c, and C57BL/6J mice using the Qiagen QIAmp DNA Kit (Qiagen, Hilden, Germany). The DNA concentration was measured using the Qubit 2.0 Fluorometer (Life Technologies, Carlsbad, CA, USA); and DNA purity and integrity were checked using a Nanodrop and by pulsed field gel analysis, respectively; and 50 pM DNA concentrations were used for LR sequencing of each strain. Pacific Biosciences (PacBio, Menlo Park, CA) LR SMRTbell libraries (~20kb) were prepared using Blue Pippin size selection according to the manufacturer’s protocol, and LR sequencing was performed using the PacBio Sequel II SMRT Cell system.

*Data processing and SV identification.* The PacBio raw bam format files were converted to the fastq format using the bam2fastx method using the default commands. CoNvex Gap-cost alignMents for LRs (ngmlr v0.2.7) ^12^ were used to align the raw data to the reference genome (mm10) using the aligner commends (-x pacbio -i <default> -R <default> -t 15). The alignment output files were sorted and converted into bam format files with the samtools *view* commend ^13^. SV identification for each sample was performed using Sniffles (v.1.0.12b) with parameters: -s 8 -l 50 --min_homo_af 0.7 --min_het_af 0.25 --genotype --cluster ^12^. To perform the downstream functional analysis of SVs present in each strain, Sniffles SV results were filtered to retain only those genomic positions with >50 bp changes, which were homozygous alternate calls and contained PASS tag. To identify shared and strain-specific SVs, a merged dataset was also assessed using Sniffles’s population calling pipeline (parameters: -s 8 -l 50 --min_homo_af 0.7 --genotype --cluster --Ivcf ). SVs of the merged callset were filtered if a certain SV was a transloation (breakends), or was a homozygous alternate call in the control strain C57BL6 (reference genome), or no homozygous alternate call present across the 6 new strains.

*Genomic feature annotation.* To assess the impact of a SV on a gene, the filtered merged callset for the 6 strains was annotated using the Ensemble Variant Effect Predictor (VEP) program ^14^. Based upon the intersection of a SV and a genomic region, VEP annotates the SV effect, which includes those effecting coding regions. For annotations (i.e. “downstream” or “upstream”), SV that intersected (within 1 bp) with a known regulatory region were analyzed using the BEDtools *intersect* option. To annotate variants in non-coding regions (upstream or downstream), the functional regions were assessed as using the following database information. Genomic promoter regions were downloaded from Eukaryotic Promoter Database (EPD) ^15^ and converted to BED track. Gene enhancer segments were downloaded from Vista enhancers ^16^ and filtered for the mouse genome (mm9), and the final BED tracks were converted to mm10 UCSC online (<https://genome.ucsc.edu/cgi-bin/hgLiftOver>). Mouse CpG Islands (GCI) representing the methylated regulatory region were extracted from the UCSC mm10 ftp site. Murine gene transcription start site data were retrieved from ENCODE ^17^ and DBTSS^18^. Genomic insulators within mouse genome (mm9) were downloaded from CTCFBSDB 2.0 for all cell types, and mm9 to mm10 coordinates were obtained from the UCSC genome browser. Murine gene alternative translation start sites were obtained from TISdb ^19^. Annotations for the genomic repeat elements were obtained using the *RepeatMasker* software package ^20^.

*Selection of genes expressed in brain*. Brain expression data was retrieved from the Expression Atlas database ^3^. Genes whose expression level was ≥10TRM, which is the basal level used by this database, were identified as genes that were expressed in brain.

*SR-SV analysis.* The SR SV dataset was constructed using the genome sequence of 53 inbred mouse strains ^2^. Data were realigned to GRCm38 using the SpeedSeq (v0.1.2) realign pipeline, and then SV analysis was performed using by SpeedSeq sv pipleline (Lumpy v0.2.13, CNVnator v0.4.1, SVTyper v0.7.0) with extra parameters: -d -P -g -k. The individual SV data for the 53 strains were merged, re-genotyped, copy-number annotated, and pruned using the svtools (v0.4.0) workflow. To obtain high quality SR-SV calls, we searched for SVs with the following parameters: SV size > 50 bp, homozygous rate > 94% (i.e., ≥50 strain genotypes are homozygous); at least 1 strain’s genotype is homozyous alternate; deletions < 1000 bp required the support of at least 1 split read; inversion’s MSQ (mean sample quality) > 150; QUAL >= 20.

*Manual inspection*. A visual inspection of computationally identified LR variants with Sniffles was performed using samplot (v1.0.1) (<https://github.com/ryanlayer/samplot>), which enabled low quality SVs to be removed.

*Evaluation of LD between SV and SNP alleles.* The relationship between SVs (n = 146K) identified from analysis of SR genomic sequence and previously identified SNPs (22M) in the genomes of 53 inbred mouse strains was investigated. For these analyses, a SV was treated as an individual allelic variant (even though it impacts >50 bp), which was just the same as a SNP allele. The rate of decay in the linkage disequilibrium (LD) between identified alleles was analyzed using PopLDdecay (v3.41) ^21^. Pairwise LD statistics ($r^{2}$) for SV and nearby SNPs (within ± 50 kb) were calculated using PLINK 1.90 ^22^ for all 53 inbred mouse strains. Because LD is the nonrandom correlation of relationship between alleles present at different loci, which is affected by non-random mating that does not occur among the inbred strains, LD relationships will mostly reflect the genealogy of the strains. We previously found that the 53 mouse strains with available genomic sequence could be separated into four sub-groups based upon their genome-wide genetic relatedness ^1^. To better characterize the relationship between a SV and its nearby SNPs, we also investigated whether SV alleles are in complete LD ($r^{2}$=1) with nearby SNPs (within 50 kb) within the strains in the 4 subgroups. The significance of the LD between SV and SNP alleles is obtained by calculating $Nr^{2}$, which follows a Chi-square distribution with one degree of freedom ($Nr^{2}\sim\chi_{1}^{2}$), where *N* is the number of strains. Thus, the p-values required for a SV to be perfectly linked with a SNP in the whole strain panel and or among each of the 4 sub-groups of the 53 inbred strains are $p_{53}=3.3E-13$, $p_{6}=0.014$, $p_{17}=3.7E-5$, $p_{25}=5.7E-7$ and $p_{5}=0.025$, respectively.

**Table S1. The SVs observed at each level of analysis.** SVs were identified from LR sequence data obtained from 6 inbred strains. LR sequencing produced on average mean read length of 15.65 kb and an average sequence depth of 40x. The numbers of SV identified and those remaining after quality control steps is shown for each strain. The C57BL/6 (reference) genome has very few SV because it is the reference genome that was used for this analysis; and consequently, it does not have any strain-specific SV alleles. In contrast, the 129Sv1 genome has the greatest number of strain-specific SVs.

|  | **129Sv1** | **AJ** | **BALB** | **BTBR** | **C57BL/6** | **SJL** |
| --- | --- | --- | --- | --- | --- | --- |
| Mean read length (bp) | 16939 | 15770 | 14162 | 15872 | 16716 | 14453 |
| Breath of coverage (%) | 93.57 | 96.55 | 96.61 | 95.97 | 94.04 | 96.56 |
| Depth of coverage (X) | 42.09 | 42.69 | 37.50 | 38.63 | 40.01 | 39.19 |
| # SV (total) | 68977 | 66903 | 59703 | 59933 | 8850 | 66787 |
| # SV (filtered) | 48292 | 48372 | 41415 | 41528 | 5482 | 45148 |
| Strain Specific | 9032 | 5648 | 3491 | 6018 | 0 | 8537 |

**Table S2.** The 53 inbred strains were divided into the four sub-groups based on their pattern of genome-wide allelic sharing ^1^.

| Group | Number of Strains | Strain List |
| --- | --- | --- |
| 1 | 6 | B10, C57BL6NJ, C57BL10J, C57BRcd, C57LJ, C58 |
| 2 | 17 | 129P2, 129S1, 129S5, BPL, BPN, BTBR, CEJ, ILNJ, KK, LPJ, NZB, NZO, NZW, PJ, RBF, SMJ, WSB |
| 3 | 25 | A/J, AKR, BALB, BUB, C3H, CBA, DBA, DBA1J, FVB, LGJ, MAMy, MRL, NOD, NON, NOR, NUJ, PLJ, RFJ, RHJ, RIIIS, SEA, SJL, ST, SWR, TALLYHO |
| 4 | 5 | CAST, MOLF, PWD, PWK, SPRET |

**Table S3.** (**A**) The numbers of SVs that were identified using SR sequence in 53 inbred strains and those within each of four sub-groups of strains are shown. SVs are categorized as either strain-unique (present in only one strain) or strain-shared (present in ≥2 strains). The number of SVs that are in perfect linkage disequilibrium (LD) ($r^{2}=1$) with nearby SNPs (within ± 50 kb) are shown for each strain group. The mean and median numbers of nearby SNPs that are in complete LD ($r^{2}=1$) with a SV are also shown. (**B**) The same analysis was also performed for SV within the 21,832 protein coding genes in the mouse genome. For each gene within the subset of genes that contain a SV, the average number of SVs that are in perfect linkage disequilibrium (LD) ($r^{2}=1$) with nearby SNPs (within ± 50 kb) and the average number of SNPs that are completely linked with a SV are shown for each group of strains.

**A**

|  | SV Type | # SVs | # SVs in LD with nearby SNPs | Median # SNPs in LD with nearby SV | Mean # SNPs in LD with nearby SV |
| --- | --- | --- | --- | --- | --- |
| 53 strains | unique | 65992 | 1956 | 2 | 4.0 |
|  | shared | 79828 | 32748 | 4 | 4.5 |
| Sub-group1  (6 strains) | unique | 4733 | 4199 | 18 | 16.7 |
|  | shared | 6710 | 6339 | 18 | 16.7 |
| Sub-group 2  (17 strains) | unique | 15682 | 11848 | 12 | 10.8 |
|  | shared | 30641 | 25640 | 10 | 9.5 |
| Sub-group 3  (25 strains) | unique | 11561 | 9968 | 14 | 12.1 |
|  | shared | 28683 | 24154 | 11 | 10.5 |
| Sub-group 4  (5 strains) | unique | 79591 | 67095 | 5 | 6.0 |
|  | shared | 34495 | 24566 | 5 | 5.5 |

**B**

|  | # of genes with a SV | Mean # of SVs per gene | Mean # SV in LD with nearby SV | Mean # of SNPs in LD with SV |
| --- | --- | --- | --- | --- |
| All 53 strains | 11449 | 4.8 | 1.0 | 4.5 |
| Sub-group1  (6 strains) | 1937 | 2.1 | 1.9 | 16.6 |
| Sub-group 2  (17 strains) | 6268 | 2.6 | 2.1 | 10.0 |
| Sub-group 3  (25 strains) | 5725 | 2.5 | 2.1 | 10.9 |
| Sub-group 4  (5 strains) | 10679 | 4.1 | 3.2 | 5.6 |

**Table S4. BTBR-unique SVs.** The gene name, functional consequence, location, type, and size of BTBR-unique SVs that disrupt exons are shown.

| Gene | Consequence | Location | type | size (bp) |
| --- | --- | --- | --- | --- |
| *Zfp987* | coding_sequence | 4:146124773-146124814 | insertion | 84 |
| *Zfp872* | inframe_deletion | 9:22200462-22200602 | deletion | 134 |
| *Scn11a* | coding_sequence | 9:119778919-119778920 | insertion | 190 |
| *Hmmr* | coding_sequence | 11:40710209-40710210 | insertion | 193 |
| *Trgc3* | coding_sequence | 13:19261724-19262719 | deletion | 996 |
| *Gm17087* | transcript_ablation | 17:8560388-8567457 | deletion | 7070 |
| *Abca17* | coding_sequence | 17:24286687-24289823 | deletion | 3137 |
| *Abca17* | coding_sequence | 17:24290168-24295081 | deletion | 4914 |

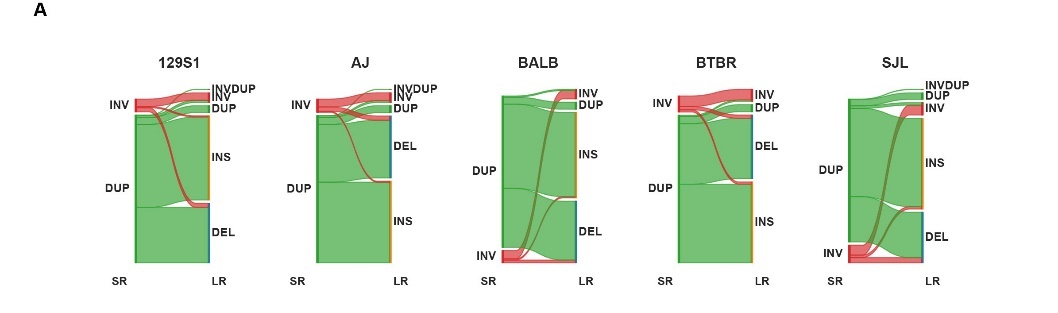

**Figure S1.** Comparison of the type of SV identified by SR and LR sequence analysis. (**A**) These Sankey diagrams indicate the number of SR-SV that were re-interpreted after analysis of the LR sequence data for each of the 5 inbred strains. Many SR-SV that were identified as inversions or duplications were not confirmed by analysis of the LR sequence data. On average, only 5% of the SV that were identified as duplications by the SR sequence analysis were not altered after LR sequence analysis, while 61.3% of the inversions were not altered.

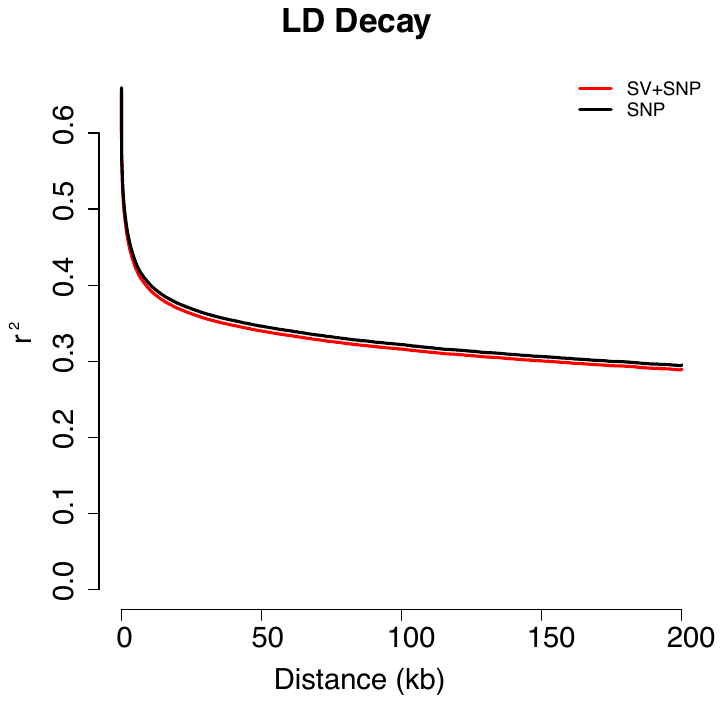

**Figure S2.** A graph of linkage disequilibrium (LD) decay for 53 inbred mouse strains using alleles identified in SNPs alone (black) or those in both SVs and SNPs (red). The half decay distance for SNPs and SVs (30-38 kb) and for SNPs alone (31-40 kb) are similar.

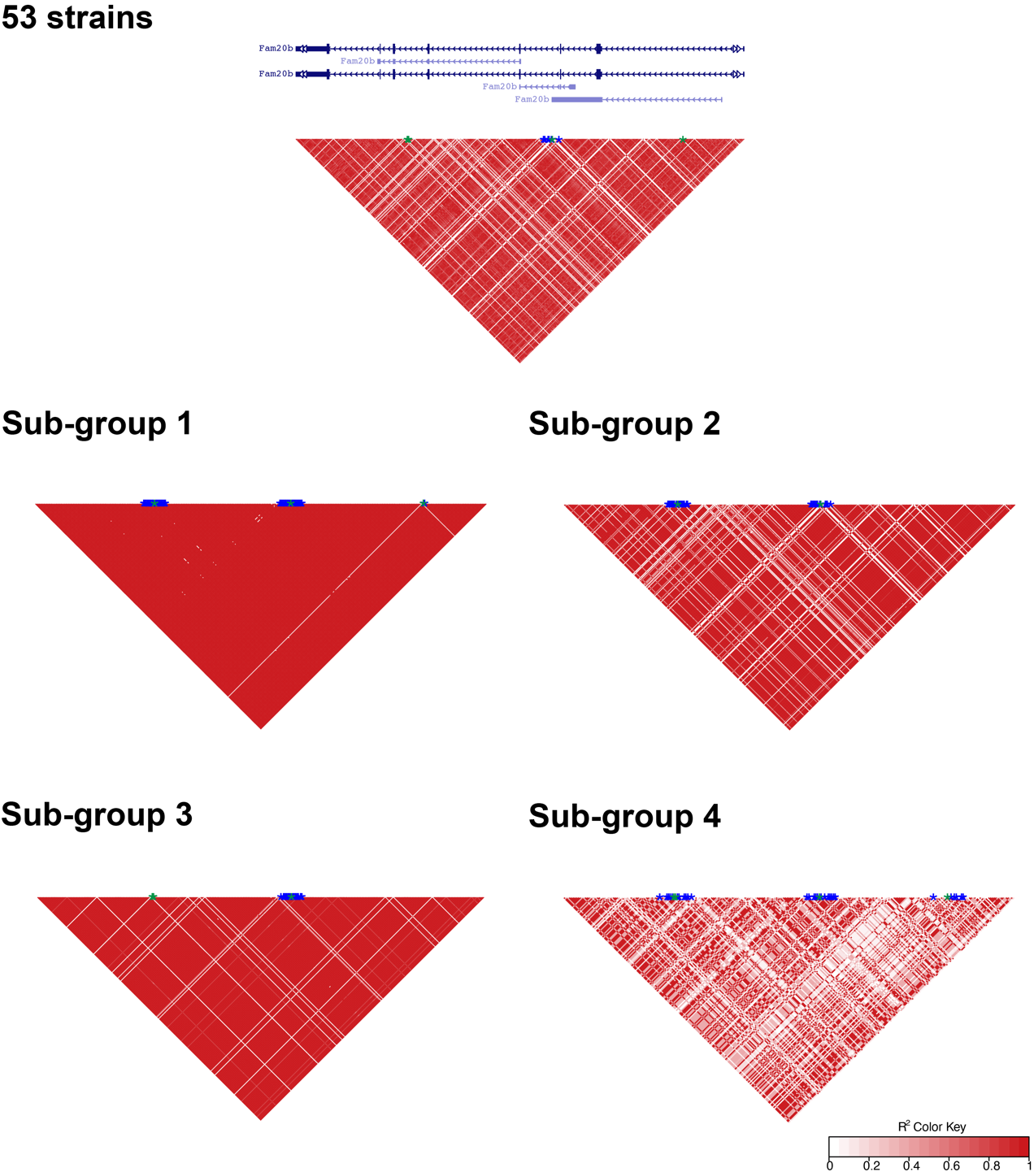

**Figure S3.** An LD plot characterizing the relationship between SVs and SNPs within the 40.5 KB *Fam20b* genomic region. *Top*: This diagram shows the locations of *Fam20b* exons with arrowheads indicating the direction of transcription, which was retrieved from UCSC Genome Browser GRCm38/mm10 mouse assembly. *Bottom*: Pairwise LD ($r^{2}$) of SVs (each indicated by a green star) and SNPs within the *Fam20b* genomic region for the 53-strain panel and for the 4 sub-groups of strains identified by population structure analysis. The calculated LD for any pair of allelic variants is indicated by the color of the region within reverse triangle, which is shown in the key below. SNPs that are in the complete LD ($r^{2}=1$) with SVs at nearby sites are indicated by a blue star. Thus, *Fam20b* has 4 SVs among the 53-strains, but only one is in complete LD with 6 nearby SNPs. The green stars for two SVs on the left partially overlap because of their proximity (chr1:156685046 and chr1:156685155), and nearby SNPs were not identified. Two SVs are present in the group 2 strains, and they are in complete LD with nearby SNP alleles. One of the 3 SVs present in the group 3 strains is completely linked with 16 nearby SNP alleles among the group 3 strains. In contrast, all 4 SVs present in group 4 strains are in complete LD with an average of 7.5 nearby SNPs.

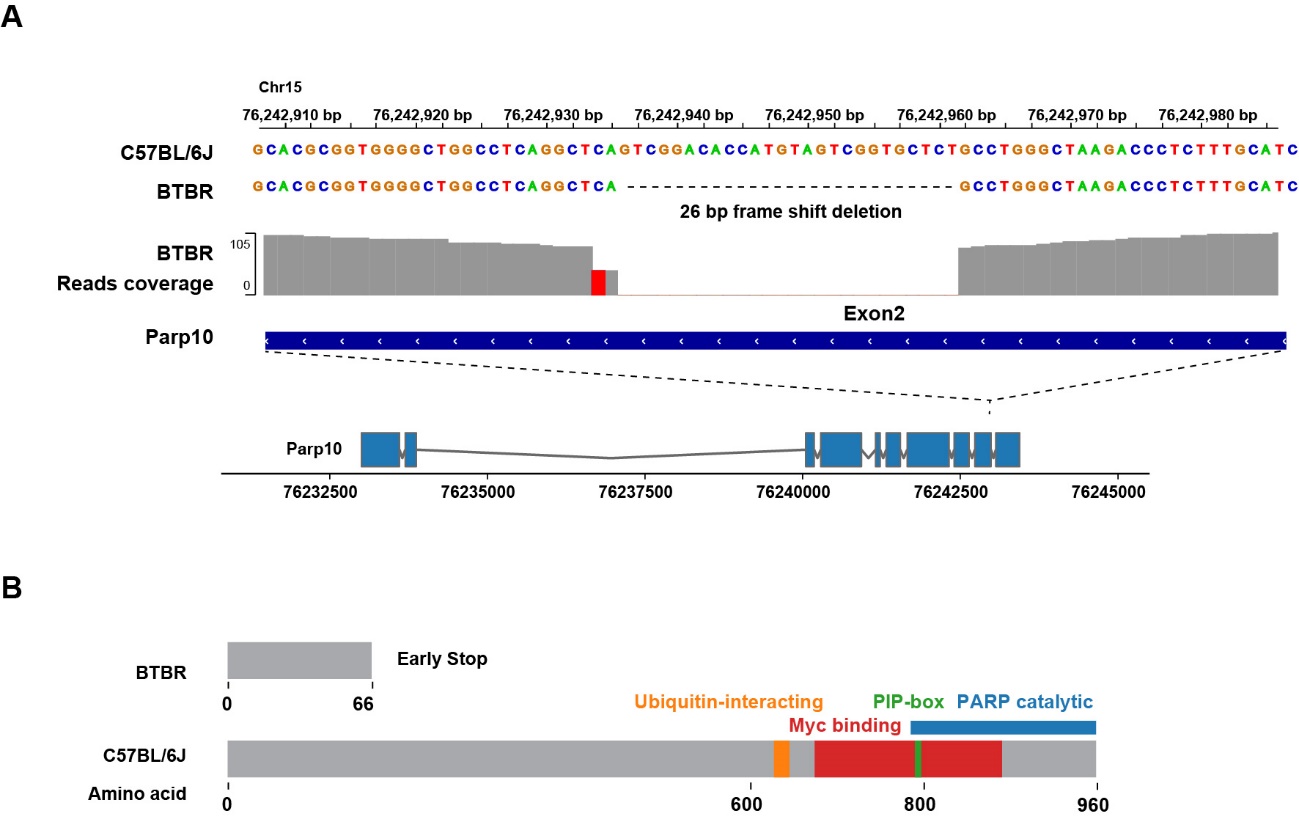

**Figure S4**. BTBR mice produce a non-functional Parp10 protein. **A**) BTBR had a 26 bp deletion in exon 2 of *Parp10*, which is not present in 52 other strains with available genomic sequence. **B**) While the full length Parp10 protein has 960 amino acids; the 26 bp frameshift deletion generates a termination codon after amino acid 66 of the BTBR Parp10 protein. The truncated BTBR protein lacks the catalytic, Myc binding, and ubiquitin-interacting domains that are essential for the role that Parp10 plays in neurodevelopment.
